# Learning to Control Complex Rehabilitation Robot Using High-Dimensional Interfaces

**DOI:** 10.1101/2022.03.07.483341

**Authors:** Jongmin M. Lee, Temesgen Gebrekristos, Dalia De Santis, Mahdieh Nejati-Javaremi, Deepak Gopinath, Biraj Parikh, Ferdinando A. Mussa-Ivaldi, Brenna D. Argall

## Abstract

Upper body function is lost when injuries are sustained to the cervical spinal cord. Assistive machines can support the loss in upper body motor function. To regain functionality at the level of performing activities of daily living (e.g., self-feeding), though, assistive machines need to be able to operate in high dimensions. This means there is a need for interfaces with the capability to match high-dimensional operation. The body-machine interface provides this capability and has shown to be a suitable interface even for individuals with limited mobility. This is because it can take advantage of people’s available residual body movements. Previous studies using this interface have only shown that the interface can control low-dimensional assistive machines. In this pilot study, we demonstrate the interface can scale to high-dimensional robots, can be learned to control a 7-dimensional assistive robotic arm, to perform complex reaching and functional tasks, by an uninjured population. We also share results from various analyses that hint at learning, even when performance is extremely low. Decoupling intrinsic correlations between robot control dimensions seem to be a factor in learning—that is, proficiency in activating each control dimension independently may contribute to learning and skill acquisition of high-dimensional robot control. In addition, we show that learning to control the robot and learning to perform complex movement tasks can occur simultaneously.

## I. Introduction

Functionality to the upper body is forfeited when individuals sustain injuries to the cervical spinal cord. How much is forfeited and how much availability of muscle groups remains to the central nervous system (CNS) can vary drastically depending on the severity of injury, the time since the onset of injury, and the scale in which the reorganization in the brain can occur [1]. Assistive machines have shown to be effective at providing support to people with loss of motor function. For instance, wheelchairs allow patients with cervical spinal cord injury (cSCI) to navigate the physical world using simple control interfaces such as sip-and-puff and head-operated joysticks. To achieve the level of functionality and ability to control assistive machines to perform activities of daily living (e.g., self-feeding), though— without assistance from caregivers—demands machines and interfaces with the capability to operate at higher degrees-of-freedom.

Wearable sensor technologies have been used to interface a person’s body movements to control machines such as assistive and rehabilitation devices and robots [2], [3]. In general, this interface is known as the body-machine interface (BoMI) [2]; it is non-invasive, capitalizes on a patient’s residual availability of body movements, and can adapt at the interface-level [4], [5]. A common strategy to control a robot using the interface is to engineer a decoder that is designed to map the high-dimensional body movements to a lower-dimensional robot control signal space. Whenever the intrinsic dimension of the body motions are higher than the machine to be controlled, dimensionality reduction techniques, such as principal component analysis (PCA) [6] or autoencoders [7], can be used to implement efficient simultaneous and continuous control of lower-dimensional devices [8], [9], [10], [11]. However, the design and the operation of such interfaces become challenging when redundancy of the body signals is reduced due to pathological conditions that impact mobility or when controlling complex multi-articulated robotic machines [12], [13]. In addition, it has yet to be shown that the BoMI is (1) scalable to high-dimensional assistive robots, such as multi-jointed robotic arms and (2) learnable by uninjured and injured populations. Moreover, if it is learnable, it is unclear (3) how learning is achieved and (4) whether reaching and functional tasks can be learned alongside learning to control.

In this paper, we demonstrate that the BoMI is (1) scalable to a 7-DoF assistive robotic arm and (2) learnable, with some constraints, by uninjured individuals. We also show results from analyses that start to address (3) how learning can occur, even when task performance is low. We hypothesize that uncorrelating dimensional couplings is important to performance and learning and share our current observations that seems to show signs that this holds true. Lastly, we show that complex movement tasks and robot control can be learned simultaneously.

We provide some background material in Section II, describe the experimental methods in Section III, share our results in Section IV, and discuss our findings and their implications in Section V.

## II. Related Work

Restoring functionality to individuals with cSCI can mean restoring independence and quality of life. Assistive machines can support people with their loss in functionality and can act as a replacement to their loss or as an intervention for physical rehabilitation. They can range from simple machines such as wheelchairs to more complex machines, such as robotic arms. In general, the level of complexity varies with the number of degrees-of-freedom (DoFs) the machine can operate. Assistive devices that require higher degree-of-freedom control, such as robotic arms (e.g., 6 or 7 degrees-of-freedom), create problems for typical assistive interfaces. This is due to a mismatch in the number of inputs the operator has access to on the interface (1 or 2) versus the number of output dimensions necessary to control the device (6 or 7). As a result, the mismatch requires people to discretely traverse the output control dimensions via mode switching, which has shown to increase time, increase cognitive load, and decrease user satisfaction [14]. In addition, existing interfaces do not allow patients to control robots simultaneously and continuously with more than two DoFs, such as robotic arms. Moreover, these interfaces do not factor the residual movement available to the patients, which can be significantly different, even if the level of injury is diagnosed to be categorically similar [1].

There is a long history of studies that suggest that even with spinal cord injury (SCI), sensory motor practice can lead to neuroplasticity in the central nervous system [15], [16]. This can hold true for even populations who are quadriplegic or tetraplegic and suffer C3 and C4 levels of injury. Evidence for plastic changes has been extended in recent work, showing that the use of the body-machine interface (BoMI) and motor practice and exercise can result in increased shoulder strength and muscle force [17]. However, these promising benefits have yet to be shown for populations with severe levels of spinal cord injury, as well as with high-dimensional assistive robots.

Nevertheless, studies such as [17], [2] involving patients with cSCI as well as uninjured populations provide us the assurance this is a viable leap. The BoMI has also been shown to be a suitable interface to control a 6-DoF assistive robotic arm, with the help of robot autonomy [3]. Furthermore, there is rich body of literature from an adjacent community made up of brain-computer interface (BCI) researchers, where technologies, approaches, and scientific questions align greatly. Instead of trying to capture residual signals from the upper body, BCI researchers capture signals from the brain or muscles using methods that range from non-invasive methods such as electroencephalogram (EEG) and electrocardiogram (ECG) to invasive methods such as surgically implanting electrodes and directly recording from the brain [18]. For example, principal component analysis (PCA)—the dimensionality reduction technique deployed in this study—is a common algorithm used to study human movement and decoder design for brain-computer interfacing.

## III. Methods

### A. Materials

The sensor net consists of four inertial measurement unit (IMU) sensors (Yost Labs, Ohio, USA), placed bilaterally on the scapulae and upper arms and anchored to a custom shirt designed to minimize movement artifacts. The relative quaternion orientation of the four IMUs in the net (16-dimensional) is mapped to a 6-dimensional subspace using PCA. The PCA map is precomputed using data from an experienced user, performing a predefined set of movements, and this same map is used for all participants. The lower-dimensional subspace consists of 6D velocity commands—3D position (*x, y, z*) and 3D rotation (*roll, pitch, yaw*)— which are used online to control a 7-DoF JACO robotic arm (Kinova Robotics, Quebec, Canada). A GUI is displayed on a tablet to provide a visualization, for the participant, of the robot velocity control commands as well as a score for each trial.

### B. Protocol

There are three phases to the study protocol: (a) familiarization, (b) training, and (c) evaluation (Figure 3). During *familiarization*, participants are encouraged to explore and become familiar with the system on their own, with minimal constraints enforced. Both of the next phases make use of a set of ten fixed targets. *𝒢* During *training*, two categories of reaching tasks are employed: reaches from a fixed center position out to a target *g*_*i*_ *∈ 𝒢*, and sequential reaches between multiple targets *g*_*j*_ *∈ 𝒢*. The ordering of targets is random and balanced across days to avoid ordering effects, and it is identical across participants. The *evaluation* phase is split into a reaching and a functional task. In the reaching task, participants reach to five targets that comprise a 3D-star *g*_*k*_ *∈ 𝒢* in fixed succession. The functional tasks are designed to emulate four ADL tasks: (a) take a cup (upside-down) from a dish rack and place it (upright) on the table, (b) pour cereal into a bowl, (c) scoop cereal from a bowl, and (d) throw away a mask in the trash bin.

**Fig. 1:**
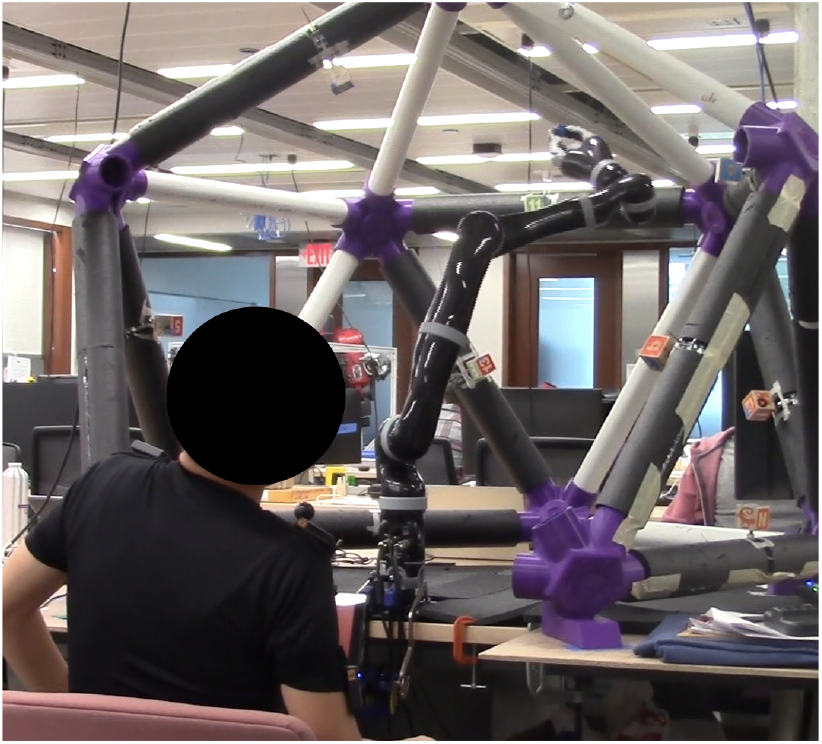
An uninjured participant attempts to control the 7-DoF robotic arm using the body-machine interface to complete a reaching task.

**Fig. 2:**
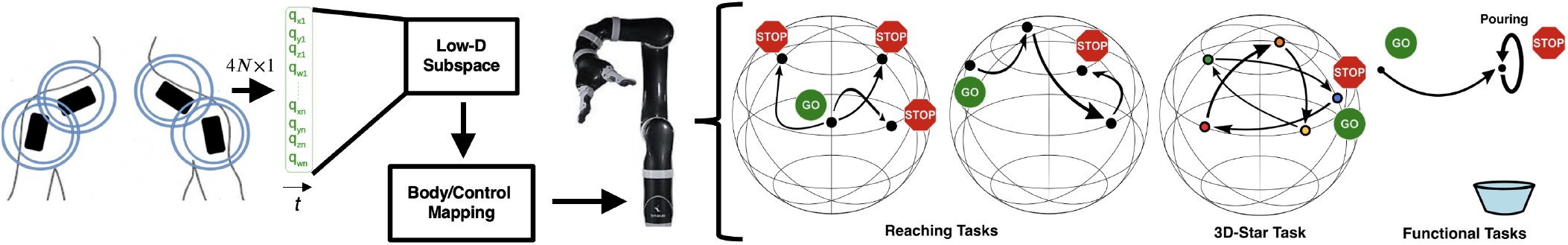
An overview of the interface-robot pipeline and the study tasks. Participants are asked to wear a sensor net on their upper bodies that control the JACO robotic arm in either task-space (3D position + 3D orientation) or joint-space (6D joint angles) to perform reaching and functional tasks. The relative quaternion orientation of the four IMUs in the net (16-dimensional) is mapped to a 6-dimensional subspace using PCA. The PCA map is precomputed using data from an experienced user, performing a predefined set of movements, and this same map is used for all participants. The lower-dimensional subspace consists of 6D velocity commands—3D position (*x, y, z*) and 3D rotation (*roll, pitch, yaw*)—which are used online to control a 7-DoF JACO robotic arm.

**Fig. 3:**
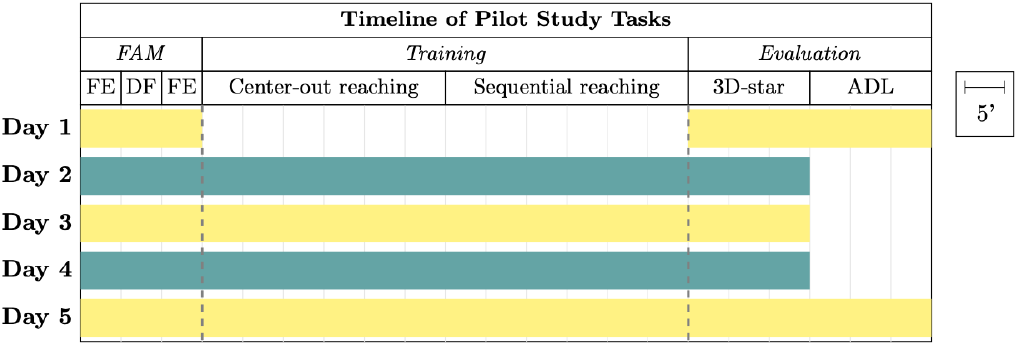
A summary of the pilot study protocol. Tasks include: (1) Free Exploration (FE); (2) DoF all-but-one freezing (DF); (3) Training, Reaching; (4) 3D-star Evaluation, Reaching; (5) Evaluation, Functional ADLs. Each row describes the various tasks participants are asked to perform on a particular day, and each column describes how often each task is performed throughout the entirety of the pilot study.

A trial ends upon successful completion or timeout. For reaching any target *g ∈ 𝒢*, success is defined within a strict positional (1.00 cm) and rotational (0.02 rad, or 1.14°) threshold, and the timeout is 90 seconds. For the functional tasks, experimenters follow codified guidelines to determine when the tasks complete and the timeout is 3 minutes. Participants are informed of the timeouts and asked to perform tasks to the best of their ability. If there is any risk of harm to the participant or the robot, study personnel intervene and teleoperate the robot to a safe position before proceeding.

### C. Participants

Each participant completes five sessions, executed on consecutive days for approximately two hours each. All sessions are conducted with the approval of the Northwestern University IRB, and all participants provide their informed consent. Ten uninjured participants from this study are reported in this paper.

## IV. Results

Figure 4 shows the results of the correlation coefficient analysis between the six decoded robot control signals, on the 3D-star evaluation task. Figure 4a shows that the group’s (*N* = 10) control signal data shares negative linear relationships in 3 of the 15 combinations of control dimensions (*𝒟* = {(1, 5), (3, 4), (4, 6)} ; *p <* 0.05). Signs of positive relationships are also shown between*𝒟* = {(3, 6), (4, 5)}, ut these combinations at a group-level are not statistically significant. Figure 4b distinguishes how correlation coefficients may differ between individuals. While the data of P_1,*lo*_ and P_2,*lo*_ displays no or very little correlation between control dimensions—especially *𝒟* = {1, 2} —P_1,*hi*_ and P_2,*hi*_ shows strong signs of correlation between dimensions, both positive and negative. More specifically, 5 of the 15 (33.3%) combinations for P_1,*hi*_; and 6 of the 15 (40%) combinations for P_2,*hi*_. Interestingly, among the dimensional combinations with statistical significance, 80% or more are correlated negatively for P_1,*hi*_ and P_2,*hi*_, which corroborates somewhat with the group analysis, where 100% (3 out of 3) of the combinations are negatively correlated.

**Fig. 4:**
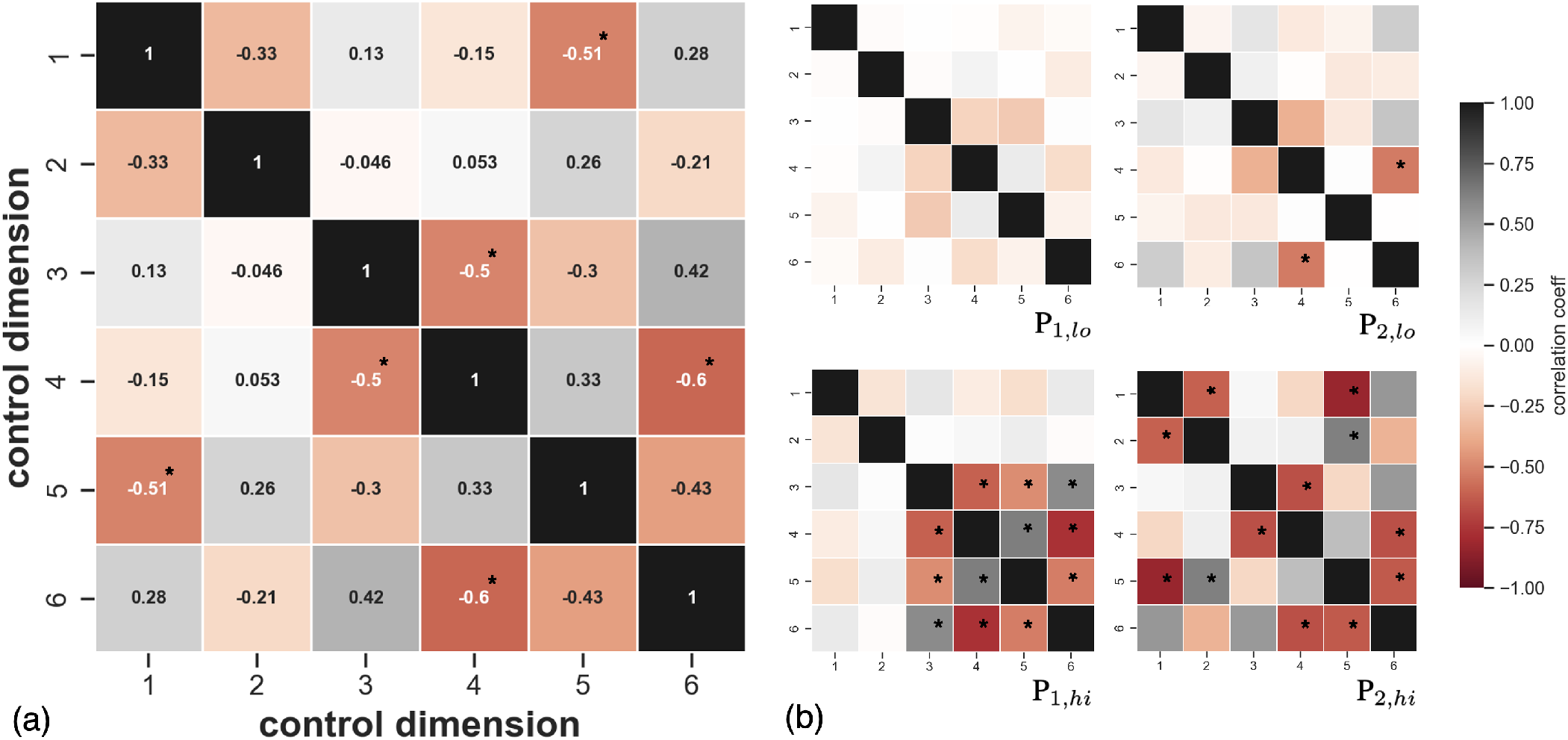
Correlation coefficients (*ρ*) between the six decoded robot control signals, collected during 3D-star evaluation task. The control signal data is represented as the magnitudes of applied robot commands. (a) Annotated group analysis; (b) analysis on selected participants, where P_1,*lo*_ and P_2,*lo*_ (top) show relatively little to no correlation between control dimensions, P_1,*hi*_ and P_2,*hi*_ show relatively high levels of correlations. Correlation coefficients range between [*−*1, +1], where *ρ* = [0, +1] implies positive linear correlation, *ρ* = [*−*1, 0] implies negative correlation, and *ρ* = 0 implies no correlation between variables. ^*∗*^*p <* 0.05

The change in variances of the magnitudes of applied robot control commands, on the 3D-star evaluation task, are shown in Figure 5. In general, there are only minor changes in variances between Days 1 and 5 in Figure 5a. Of the minor deltas, two-thirds of the control dimensions (*𝒟* = {2, 3, 5, 6}) increase, whereas only one dimension decreases. Variance of *𝒟*4 appears to change the least, as well as result in the smallest variances (0.007 compared to [0.011, 0.020]). Lastly, the spread of the variances appear to be consistent in the group. Figure 5b shows the same analysis on the selected participants. A significant decrease in variances is experienced by P_2,*lo*_, in particular on three dimensions *𝒟* = {2, 3, 5}. Moreover, the spread of variances also decreases greatly on all six control dimensions. Changes in variance for P_2,*hi*_ are major in 5 out of 6 dimensions, where variances decrease in 2 out of the 5 dimensions (*𝒟* = 1, 6) and increase in the remaining 3 dimensions (*𝒟* = {2, 3, 5}). P_1,*lo*_ and P_1,*hi*_ both do not experience much change in variance overall between first and last days. However, the two dimensions that decrease for P_1,*lo*_ (*𝒟* = 3, 5) are the two dimensions that start on Day 1 with the largest variances; P_1,*hi*_ shows a similar pattern in the fifth control dimension.

**Fig. 5:**
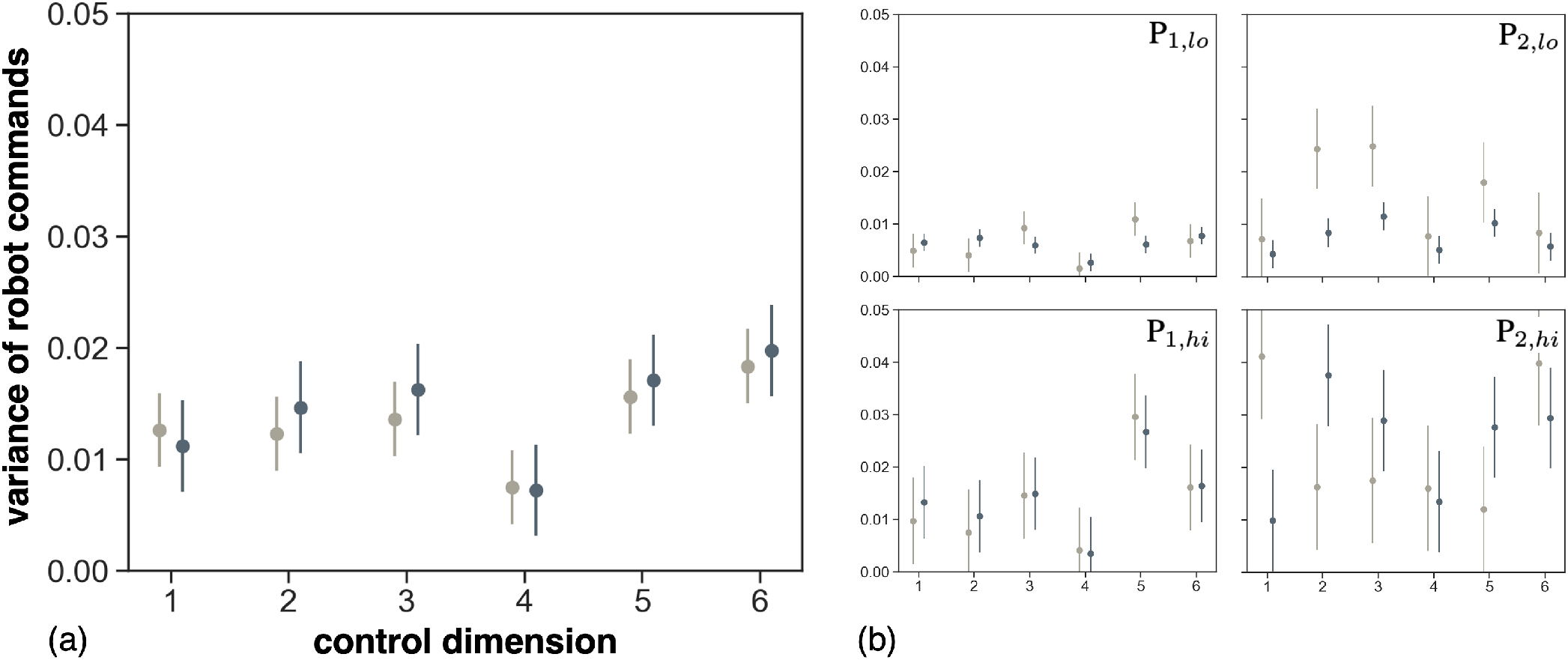
Variances of the magnitudes of decoded robot commands on the 3D-star evaluation task. (a) Group-level analysis; (b) analysis of selected participants, where selection and order is preserved from Fig. 4.

Figure 6a shows the counts of collision and intervention instances during the 3D-star task, where the selection of participants and order is preserved from Figure 4. P_1,*lo*_ experiences no interventions and only one collision on Day 3. P_2,*lo*_, however, experiences the most amount of collisions (47) among participants on Day 1, but this dramatically reduces as early as Day 2 and decreases similar to a decaying exponential. P_1,*hi*_ and P_2,*hi*_’s data show signs of similarities in the number of collisions and their flat trend across the five days. Overall, the number of collisions are relatively small, ranging from [0, 3], among the selected participants. P_2,*lo*_’s interventions occur early in the week on Days 1-3, whereas P_1,*hi*_ and P_2,*hi*_ experience interventions in the latter half of the week, on Days 4 and 5 and on Day 4, respectively. However, while the protocol has strict policies on when study personnel should intervene, there remains some level of subjectivity; therefore, this is only a small observation.

**Fig. 6:**
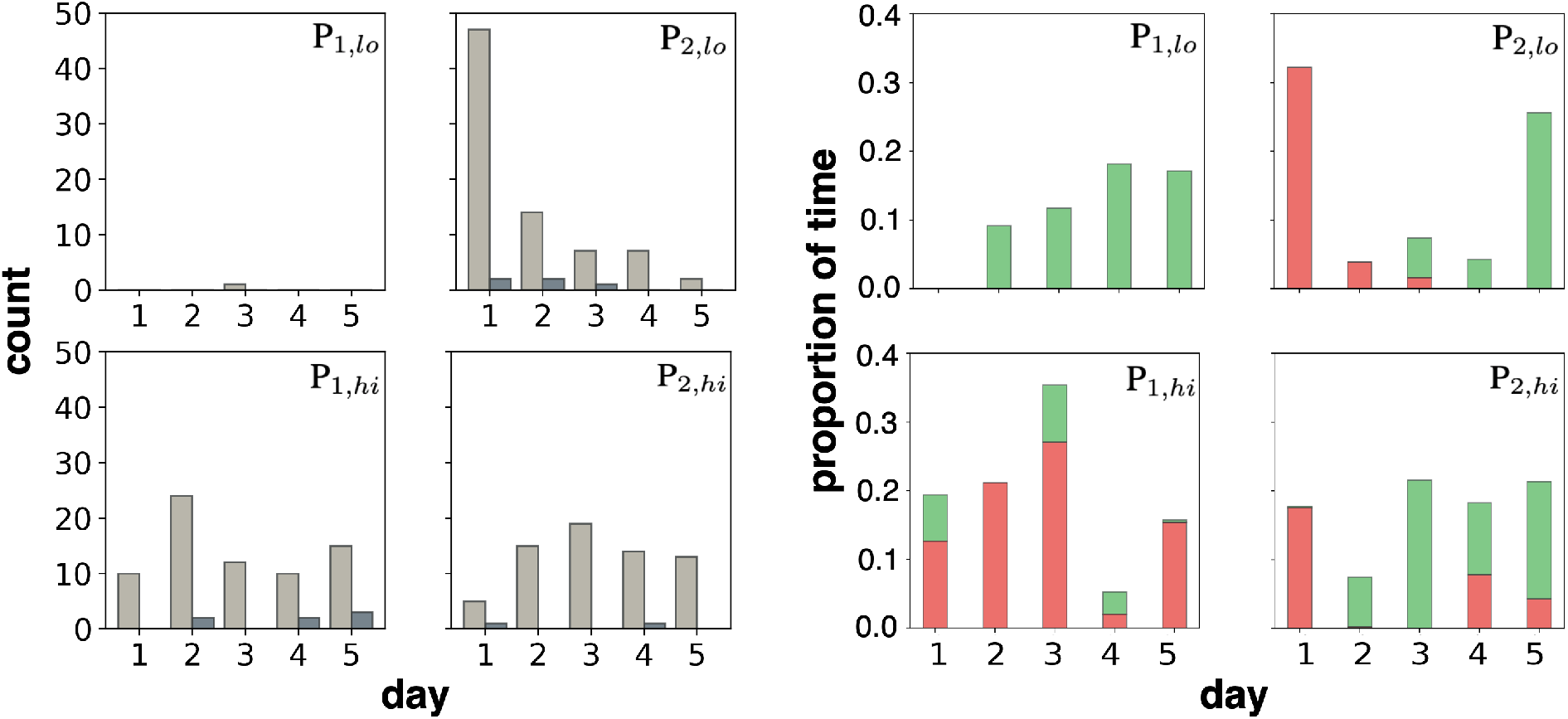
(a) Counts of collision (light) and intervention (dark) in the 3D-star evaluation task across days. (b) Proportion of time the robot endpoint spends proximal or remote, relative to reaching targets on the 3D-star evaluation task. Proximal zone (green) is within 10% of reaching distance, and remote zone (red) is outside of 100% of reaching distance. Selected participants and order is preserved from Fig. 4.

The study workspace is broken down into three distinct zones, with respect to a given target, in the analysis in Figure 6b. Especially because there were *zero* successful reaching trials, we instead demarcate three zones at 10% and 100% of the total reaching distance needed in position and orientation: (1) proximal (green), (2), peripheral (grey), and (3) remote (red) zones. Among the selected participants, P_1,*lo*_ spends no time in the remote zone (farther than 100% of the reaching distance) and increases the the time spent in the proximal zone (within 10% of the reaching distance) over the five days. P_1,*hi*_, on the other hand, spends significantly more time in the remote zone compared to the time spent in the proximal zone, spending the most amount of time in this zone on Day 3 (27%). On Days 4 and 5, though, time spent in the remote zone dramatically reduces, showing some signs of learning. P_2,*lo*_ and P_2,*hi*_ both start the week spending much of their time in the remote zone and very little in the proximal zone. By the second half of the week, they decrease their time spent in the remote zone while increasing the time spent in the proximal zone.

## V. Discussion

In this pilot study, we examined whether the body-machine interface is scalable to a 7-DoF assistive robotic arm, able to be learned by an uninjured population, even when task performance can be low, and complex tasks can be learned in parallel with learning to control. Our results suggest that uncorrelating dimensional couplings, in the control of complex robots using high-dimensional interfaces, may be involved during learning as people try to gain proficiency. They also suggest that learning to control and learning to complex tasks can occur simultaneously, rather than having to be learned serially.

The BoMI has been shown to be an effective interface to control assistive and rehabilitation devices and robots [2], [3], drones [19], and quadcopters [20]. Especially in the domain of assistive and rehabilitation machines, though, previous demonstrations on physical robots have been in only one or two dimensions of control [21], [11] and not six or seven dimensions of control. Despite some undesirable properties such as control axes not being preserved in body motion space, uninjured participants were able to learn to control the robot to perform complex reaching movements and several functional tasks.

P_1,*lo*_ seems to be an exemplar to our hypothesis that uncorrelating dimensional couplings is key to performance and learning. P_1,*lo*_ keeps variances of control command magnitudes and their spreads low along dimensions on Days 1 and 5, compared to the other participants (Figure 5). While variances can be dependent on the task, based on the results from other analyses, low variances seem to be a characteristic of the 3D-star task. In addition, instances of collision and intervention are also kept extremely low (Figure 6a); in fact, on both measures, P_1,*lo*_ shows the least amount of collisions and interventions among all ten of the participants. P_1,*lo*_ also avoids spending any time, with the robot end-effector, beyond the distance between starting position and targets and reaches within 10% of the reaching distance on all five targets by Day 5. However, it is important to note that despite outperforming all other participants in these metrics, P_1,*lo*_ was not able to complete any of the reaching tasks in the 3D-star evaluation task. We believe part of the reason is, results from Figure 6b do not account for meeting any orientation goals. Further analysis in orientation space could provide better context. An alternative explanation could be there are other aspects to learning than isolating robot control dimensions. This learning strategy is actually counterintuitive, because, in general, isolating movements along dimensions is inefficient (e.g., manhattan distance versus cartesian distance; cerebellar patients versus healthy [22]), and it is not how the human brain chooses to make voluntary movements with our anthropomorphic limbs. Uncorrelating control dimensions here is what we posit as critical to learning robot control, not to make efficient movements. Therefore, it is perhaps necessary to design additional experiments to study how individuals can learn to make efficient movements once proficiency in robot control is achieved. Another option could be to delegate movement efficiency to robot autonomy.

Similar to P_1,*lo*_, P_1,*hi*_ presents another (negative) example to our hypothesis. Variances in the last two dimensions are especially elevated, with wider spreads of variances, and do not change much between first and last days. Counts of collisions are generally high, and they do not decrease over days. We notice proportionally more time spent farther away from the targets relative to the starting distances, compared to within 10% of the distances. We also see a small increase in visits to the proximal zone and decreases in both time spent and visits to the remote zone—all positive signs in terms of learning. Interestingly, given that P_1,*hi*_ experienced less coupling in the first two control dimensions, this did not seem to necessarily result in compensatory behavior, where *D* = {1, 2} are activated more than others (not shown). Uncorrelating control dimensions was perhaps not part of this participant’s learning strategy or it was and was abandoned early on. Further analysis may show the likelihood that someone adopts this kind of learning strategy, whether implicit or explicit. In the event this is a strategy that most do not favor, but is a statistical correlate to performance and learning, prescribing learning strategies, like uncorrelating dimensions, to people may be interesting to investigate in the future.

P_2,*hi*_ could be a counterexample and was a surprise to us on measures especially assessed in Figure 6b. Even without the context of orientation measures, it is interesting to observe how this participant was able to move the robot endpoint into proximal zone 100% of trials by Day 5, spend significant time within the proximal zone on multiple days, and show desired trends in both increase in time spent in the proximal zone and decrease in time spent in the remote zone across days. This is especially considering the strong dimensional correlations P_2,*hi*_ experienced, as shown in Figure 4. This could mean that proportions of time spent within 10% or beyond 100% of the reaching distance are not strong indicators of learning. The time series analysis of endpoint distances to target in Figure 10 hints why this could be the case, based on the inconsistency and erratic patterns of this metric. More complex time series analyses may be pursued to achieve a better understanding of the relationship between dimensional coupling and performance and learning. For instance, trajectory smoothness [23] and dimensionless jerk [24] might provide a more complete picture when erratic patterns like this is observed.

It is possible that providing assistance to reduce the burden of uncorrelating dimensional couplings may facilitate learning as well and could be a promising intervention for rapidly teaching high-dimensional robot control. This especially holds true for individuals similar to P_1,*hi*_ and P_2,*hi*_ who are not able to implicitly learn to separate the robot control signals consistently. In addition, while PCA has many attractive properties, such as linearity, when designing a decoder, there were signs that the decoding map (more specifically, the covariance) had dimensional properties that were intrinsically coupled (not shared). In other words, identifying those dimensional combinations, a priori, that are intrinsically coupled may provide strategic insights on how to provide assistance. Targeting these combinations early on could speed the learning process.

Finally, we acknowledge the importance of rethinking our study design such that tasks are achievable. However, if the long-term aim is to provide a path for patients with cSCI to rehabilitate and restore motor function via controlling the robotic arm, then the solution to this issue is not simple. We can perhaps add simpler evaluations that, for instance, tests a participant to access control dimensions on demand. But, the other challenge or limitation is time, because we cannot expect patients to quickly adapt to longer training times (over 2 hours a day). We suspect the design needs a delicate balance between increasing the likelihood of task success versus time—and being vigilant about what is required to achieve the longer-term aim.

## ACKNOWLEDGMENT

Research reported in this publication was supported by the NIH T32 Training Program and Eunice Kennedy Shriver National Institute of Child Health & Human Development (NICHD) under the award number 2T32HD007418-26; the National Institute of Biomedical Imaging and Bioengineering (NIBIB), award number R01-EB024058; National Science Foundation (NSF), award number 2054406; National Institute on Disability, Independent Living and Rehabilitation Research (NIDILRR), award number 90REGE0005-01-00; and European Union’s Horizon 2020 Research and Innovation Program under the Marie Sklodowska-Curie, Project REBoT, award number GA-750464. The content is solely the responsibility of the authors and does not necessarily represent the official views of the National Institutes of Health.

## APPENDIX

**Fig. 7:**
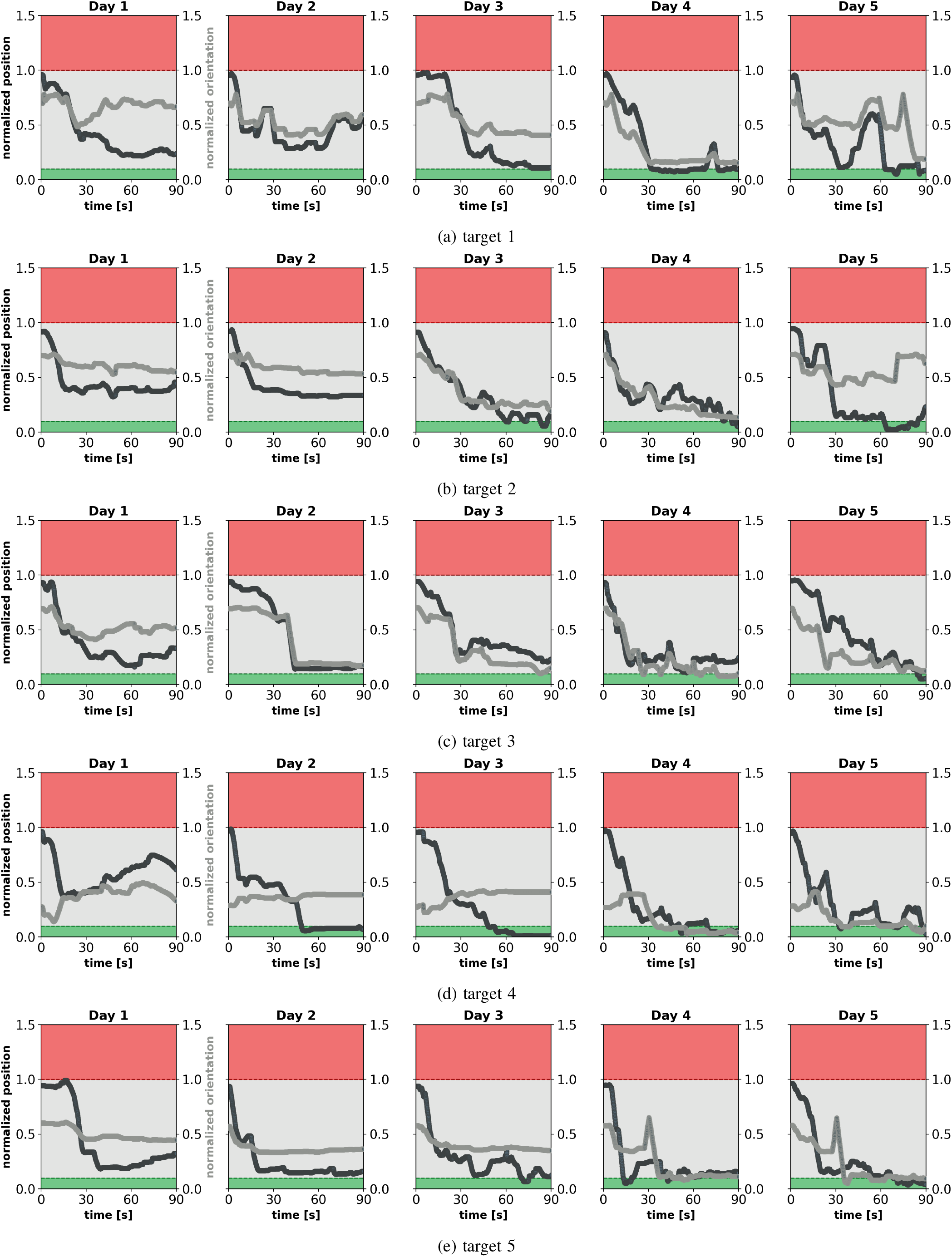
Time series of distance to target in the 3D-star evaluation task for P_1,*lo*_.

**Fig. 8:**
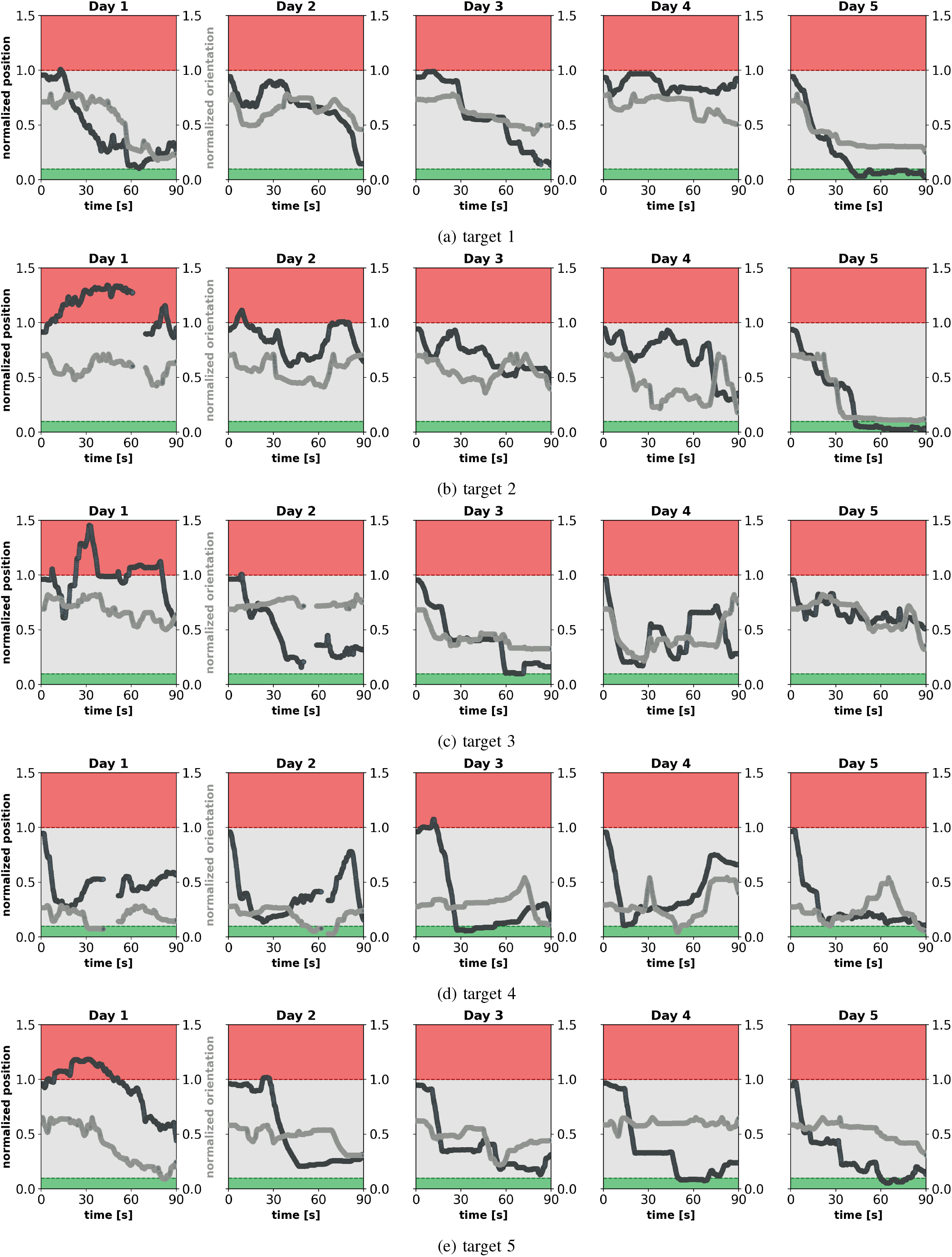
Time series of distance to target in the 3D-star evaluation task for P_2,*lo*_.

**Fig. 9:**
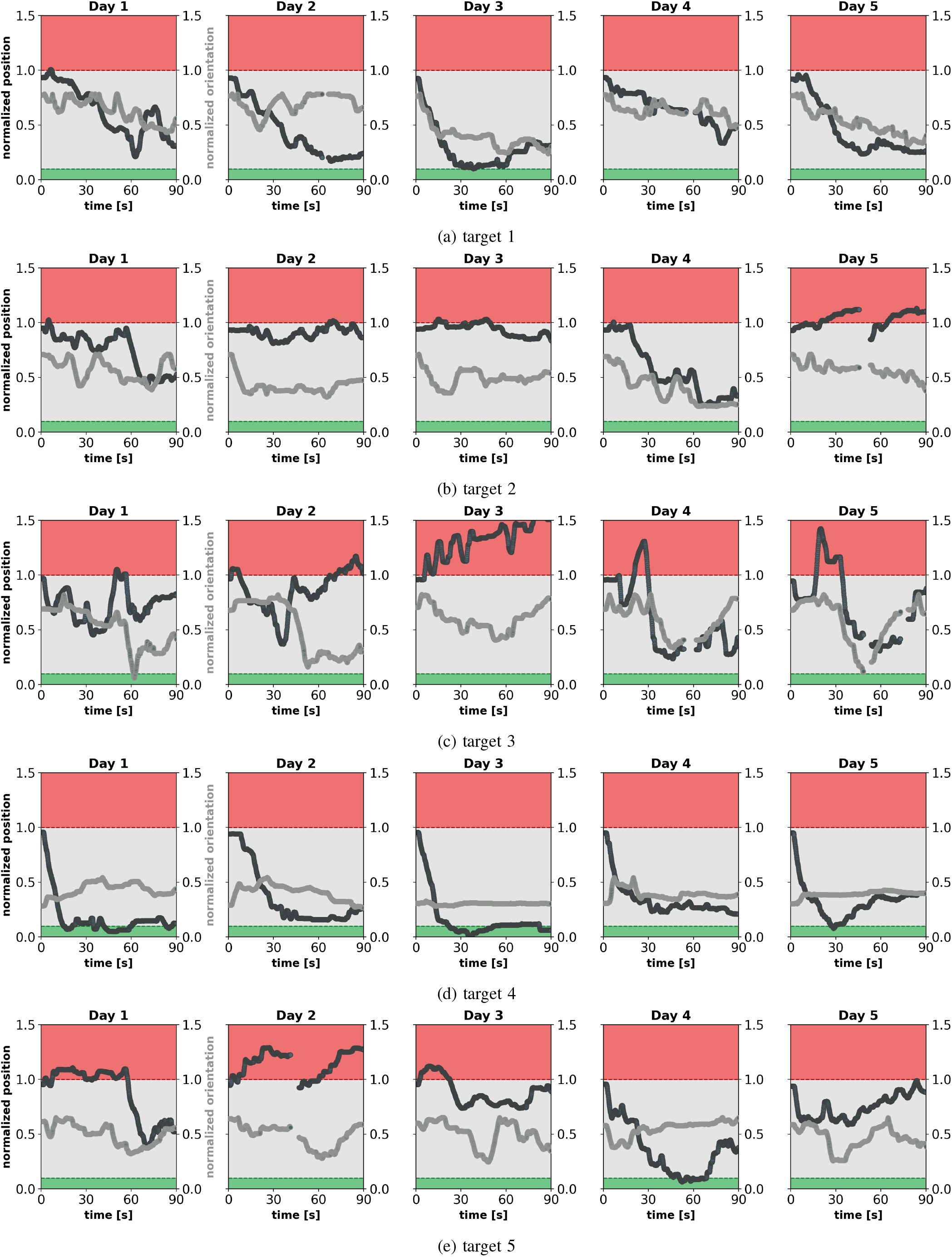
Time series of distance to target in the 3D-star evaluation task for P_1,*hi*_.

**Fig. 10:**
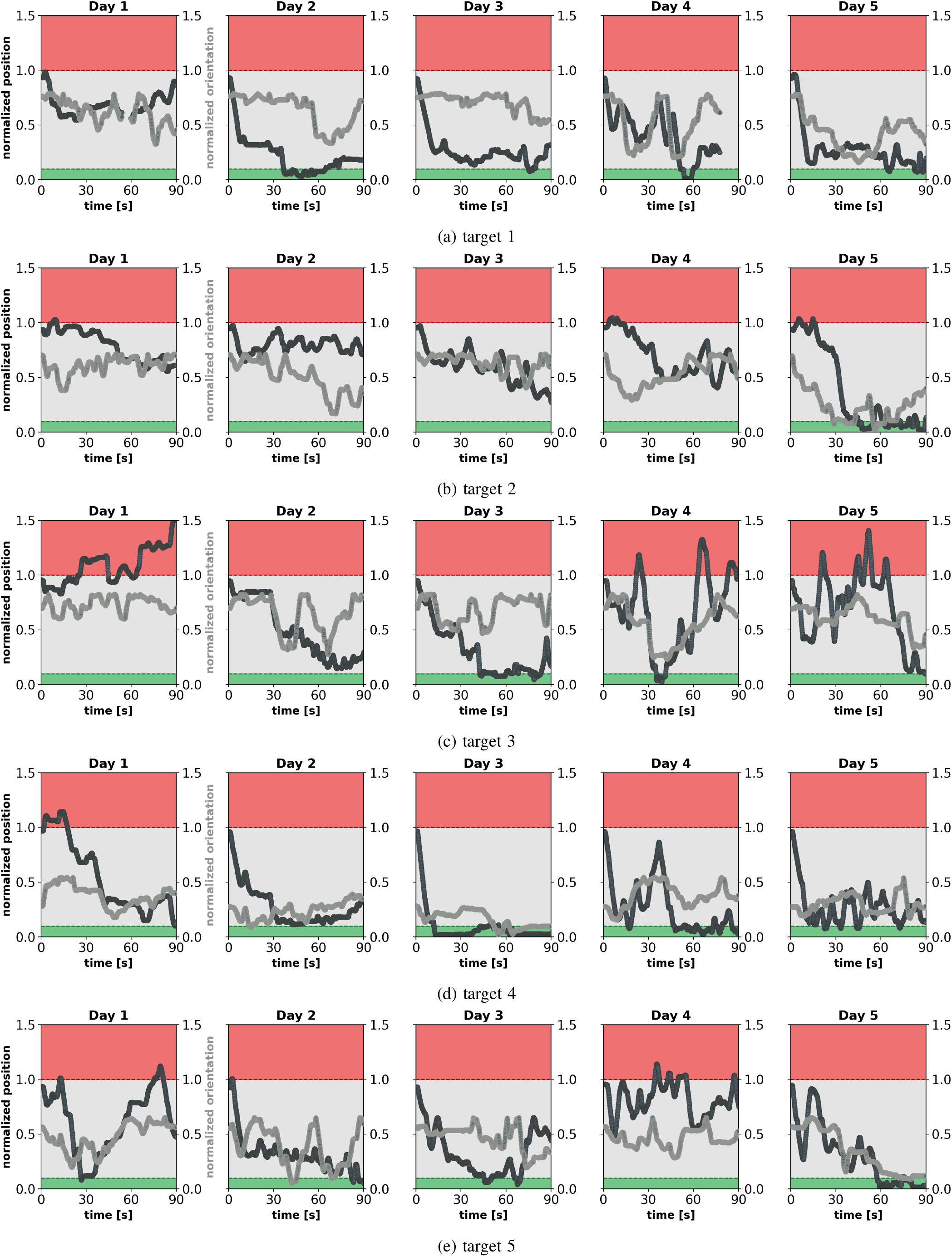
Time series of distance to target in the 3D-star evaluation task for P_2,*hi*_.

